# A changing signaling environment induces multiciliated cell trans-differentiation during developmental remodeling

**DOI:** 10.1101/2020.04.16.045401

**Authors:** Alexia Tasca, Martin Helmstädter, Magdalena Brislinger, Maximilian Haas, Peter Walentek

## Abstract

Multiciliated cells (MCCs) are extremely highly-differentiated, presenting >100 cilia and basal bodies. We analyzed how MCCs are lost from the airway-like *Xenopus* embryonic epidermis during developmental tissue remodeling. We found that some MCCs undergo apoptosis, but that the majority trans-differentiate into secretory cells. Trans-differentiation involves loss of ciliary gene expression, cilia retraction and lysosomal degradation. Apoptosis and trans-differentiation are both induced by a changing signaling environment through Notch, Jak/STAT, Thyroid hormone and mTOR signaling, and trans-differentiation can be inhibited by Rapamycin. This demonstrates that even cells with extreme differentiation features can undergo direct fate conversion. Our data further suggest that the reactivation of this developmental mechanism in adults can drive tissue remodeling in human chronic airway disease, a paradigm resembling cancer formation and progression.

## Introduction

The fate of terminally-differentiated cells appears fixed, but direct fate change through reprogramming techniques or trans-differentiation in regeneration and pathogenesis demonstrate that cell identity is more flexible than previously thought (Merrell and Stanger, 2016; Wells and Watt, 2018). This suggests that maintaining cell identity is not an intrinsic feature of differentiation but relies on extrinsic cues from the environment. While it is accepted that specialized cells can be induced from more generic types, e.g. fibroblasts or stem cells (Soldner and Jaenisch, 2018), it is debated to which degree conversion is possible in cells that have generated highly specialized morphological features.

Multiciliated cells (MCCs) are the only cells that form >100 motile cilia and basal bodies, i.e. modified centrioles generated through deuterosome-mediated amplification (Klos Dehring et al., 2013). MCCs are also subject to a permanent cell cycle block due to the deployment of centrioles to ciliogenesis, mutually exclusive with cell division (Izawa et al., 2015). Hence, it is widely considered impossible for MCCs to undergo fate change during normal development and in regeneration. In mucociliary epithelia, such as the airway epithelium or the embryonic epidermis of *Xenopus* tadpoles, correct balance between MCCs and secretory cells provides the functional basis for removal of particles and pathogens to prevent infections and to maintain organismal oxygenation (Walentek and Quigley, 2017). Mucociliary epithelial remodeling and MCC loss are observed in human chronic lung disease as well as during metamorphosis in *Xenopus*, however, it remains unresolved how and why MCCs are lost in various conditions (Hogan et al., 2014).

We studied the process of MCC loss in the developing epidermis of *Xenopus* and found that MCCs are first locally removed by lateral line-dependent apoptosis, but later globally removed through trans-differentiation into a mucus-secretory cell type. Both processes are induced by a changing Notch signaling environment and modulated by Jak/STAT, Thyroid hormone and mTOR signals. We show that MCC trans-differentiation occurs in a non-pathogenic state, involves coordinated cilia retraction, cytoskeletal rearrangement, and lysosomal elimination of ciliary and basal body material. Furthermore, Rapamycin inhibits trans-differentiation, suggesting a positive role for autophagy in the maintenance of MCCs.

## Results

Like the mammalian airway epithelium, the *Xenopus* embryonic epidermis is composed of MCCs, ionocytes, mucus-secreting cells (Goblet and SSCs), and sub-epithelial basal stem cells (Haas et al., 2019; Walentek and Quigley, 2017). Once established by stage (st.) 32, cell type composition is generally invariable between individuals (Haas et al., 2019). Using scanning electron microscopy (SEM), we found that as development proceeded, MCCs were strongly reduced at st. 45 and largely lost by st. 47, while ionocytes, SSCs and Goblet cells persisted (**Fig. 1A**).

**Fig. 1.**
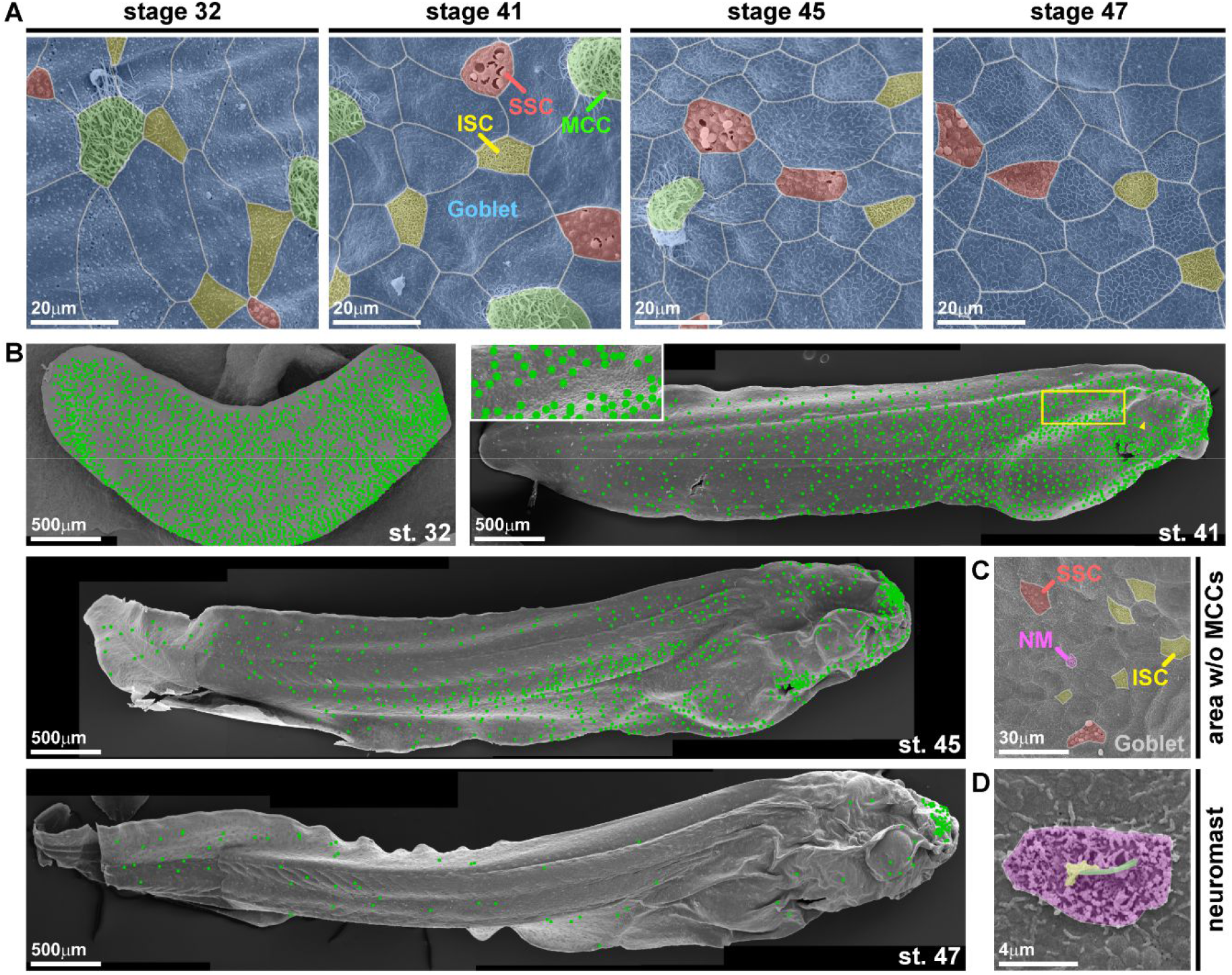
Characterization of epidermal MCC loss during *Xenopus* development. **(A-D)** Pseudo-colored scanning electron micrographs from developmental stages (st.) 32 through 47. Cf. **Fig. S1** for non-colored images. **(A)** Analysis of the changing composition of mucociliary cell types in the epidermis shows progressive loss of MCCs (green), while ionocytes (ISCs, yellow), small secretory cells (SSCs, red) and mucus-secreting Goblet cells (blue) remain present. **(B)** Analysis of MCC-loss patterns on whole tadpoles. MCCs are marked by green dots. Images were reconstructed from multiple individual micrographs. N = 7, st. 32; 6, st. 40/41; 3, st. 45; 3, st. 47. Box indicates magnified area. Arrowhead indicates MCC loss ventral to the eye. **(C)** Magnification of skin area devoid of MCCs at st. 45 reveals presence of lateral line neuromast (NM, purple). **(D)** Magnified image of a neuromast. Kinocilium (green), stereocilia (yellow), support cells (purple).

We first detected local loss of MCCs at st. 41 in areas where lateral line neuromasts (NMs) (Winklbauer, 1989) emerge by st. 45 (**Fig. 1B-D** and **Fig. S1**). The lateral line senses external fluid flows for orientation, and MCCs produce fluid flow by coordinated beating of motile cilia. Thus, we wondered if MCCs are removed to allow for lateral line function. We used correlative light and electron microscopy, immunofluorescent staining for Acetylated-α-tubulin (Ac.-α-tubulin), and a transgenic NM marker (p27∷GFP) (Carruthers et al., 2003; Rubbini et al., 2015) to investigate MCC loss and NM emergence. We confirmed that p27∷GFP is expressed in *Xenopus* NMs and that MCC loss is initially restricted to GFP(+) areas (**Fig. S2A-D**).

We also transplanted the lateral line primordium (Winklbauer, 1989) from membrane-RFP (mRFP) injected donors into hosts in which MCCs were marked by GFP expression (αtub∷GFP). This revealed MCC presence above the lateral line leading edge, but loss at positions where NMs have emerged (**Fig. 2A** and **Fig. S3A,B**). Furthermore, we found MCCs with abnormal morphology in those areas by SEM, suggesting MCC shedding and apoptosis (**Fig. S2E-G**).

**Fig. 2.**
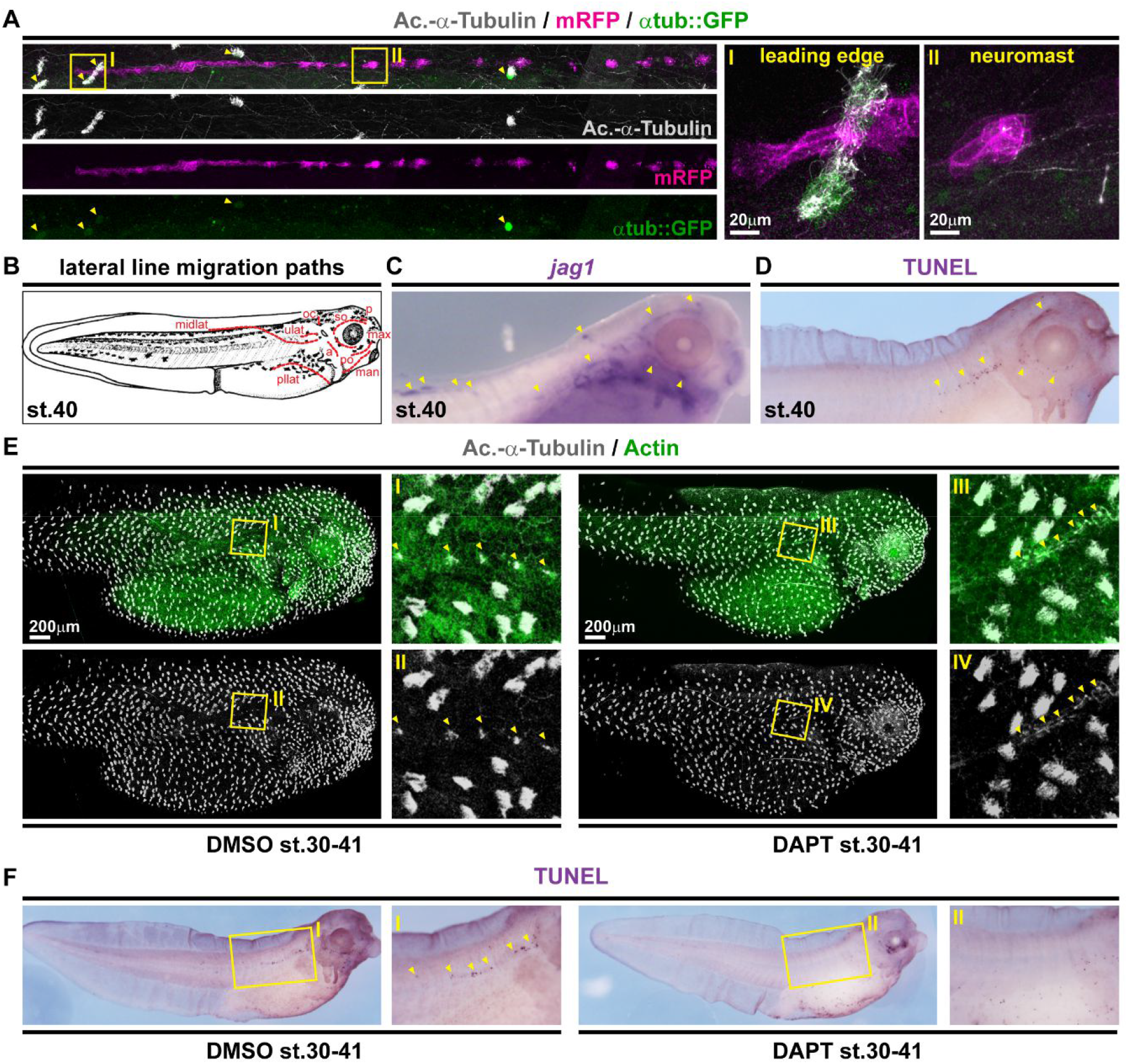
Local loss of MCCs is induced by Notch signaling derived from the emerging lateral line. **(A)** Confocal micrographs of transplanted lateral line primordium and neuromasts (marked by membrane-RFP, mRFP) reveals loss of MCCs (marked by α-tubulin promotor-driven GFP [αtub∷GFP, green] and Acetylated-α-Tubulin [Ac.-α-Tubulin, grey] antibody staining) in areas where neuromasts have emerged. Yellow arrowheads indicate GFP(+) MCCs. Cf. **Fig. S3 A,B**. Images were reconstructed from multiple individual micrographs. **(B)** Schematic representation of lateral line migration patterns in st. 40 *Xenopus* tadpoles. **(C)***In situ* hybridization shows *jagged1* (*jag1*, purple) expression in the lateral line primordium and in neuromasts. **(D)** TUNEL staining (purple) reveals apoptotic cells clustering along the lateral line migration paths. **(E)** Notch signaling inhibition by DAPT prevents lateral line induced MCC (Ac.-α-Tubulin, grey) loss in the presence of neuromasts (yellow arrowheads). F-actin (Actin, green) was used as counterstain in confocal images. Images were reconstructed from multiple individual micrographs. **(F)** Notch signaling inhibition by DAPT application prevents induction of apoptosis (TUNEL staining, purple; yellow arrowheads) by the lateral line. Magnified areas in A,D-F are indicated by yellow boxes. Cf. **Fig. S3 C,D** for additional stages related to panels C and D, and **Fig. S4** for quantification of results related to panels E and F.

The lateral line primordium and NMs are signaling centers expressing ligands from the Notch, Wnt and FGF signaling pathways that regulate morphogenesis and proliferation (Dalle Nogare and Chitnis, 2017). Notch signaling prevents MCCs during specification (Walentek and Quigley, 2017), and we tested if Notch could be activated by the lateral line to induce removal of MCCs. We analyzed expression of all Notch ligands and found that the lateral line primordium and NMs express *jagged1* (*jag1*) at high levels, and TUNEL staining demonstrated apoptosis in the epidermis along the paths of lateral line migration (Winklbauer, 1989) (**Fig. 2B-D** and **Fig. S3C,D**).

Cell type-specific apoptosis in response to Notch signaling was shown to be mediated via activation of the transcription factor Hes1 in mammalian lymphocytes (Kannan et al., 2011). We also found transient *hes1* expression in the *Xenopus* epidermis closely following the leading edge of the lateral line primordium (**Fig. S3E**). Therefore, we tested the functional relationship between Notch and MCC loss by treatment with the Notch signaling inhibitor DAPT. DAPT treatment prevented local loss of MCCs, reduced TUNEL signals as well as *hes1* expression, but without interfering with lateral line migration or *jag1* expression (**Fig. 2E,F** and **Fig. S4**). Interestingly, NMs failed to properly intercalate in DAPT treated specimens, indicating that MCCs have to be removed by Notch signaling to make space in the epidermis for NM emergence (**Fig. S4A,C**).

Extensive TUNEL staining was missing from areas further away from the lateral line (**Fig. 2D** and **Fig. S3D**) indicating an alternative mode of MCC removal there. We stained tadpoles between st. 32 and 46 for MCC cilia (Ac.-α-tubulin) and secretory cell mucus (PNA) to identify cell types, and for F-actin (Actin) to outline cell borders. Confocal microscopy revealed altered apical F-actin morphology in MCCs, which also stained positive for mucus (**Fig. 3A**). While the overall number of identifiable MCCs decreased over time, the proportion of PNA(+) MCCs increased, indicating MCC to Goblet cell trans-differentiation as an additional mechanism for MCC removal in *Xenopus* development (**Fig. 3B** and **Fig. S5D**).

**Fig. 3.**
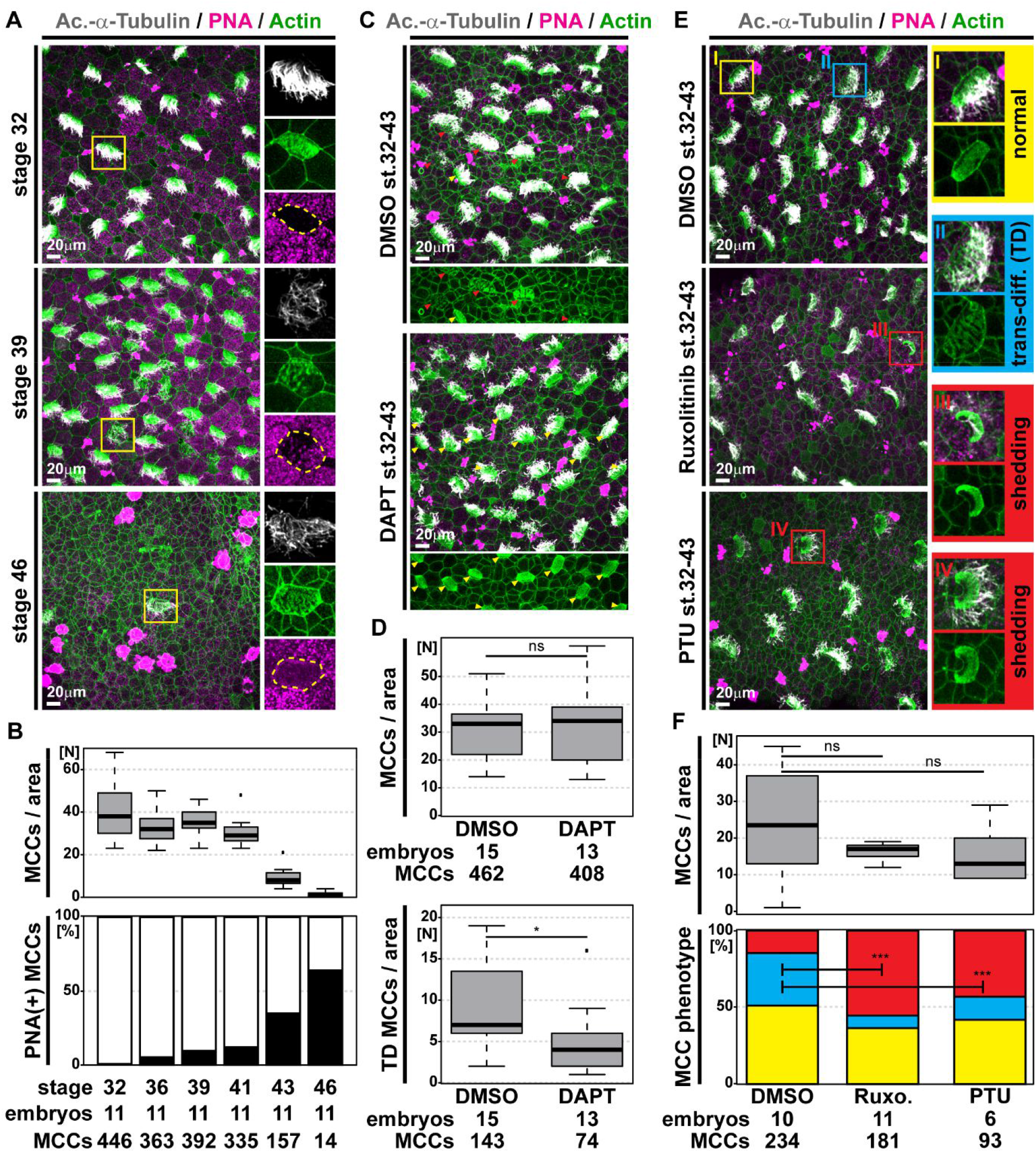
Global loss of MCCs through trans-differentiation into Goblet cells is regulated by Notch, Jak/STAT and Thyroid hormone signaling. **(A,C,E)** Confocal micrographs of the epidermis stained for MCCs (Ac.-α-Tubulin, grey), F-actin (Actin, green) and mucus (PNA staining, magenta). **(A)** Analysis of samples from st. 32 to 46 reveals MCCs with altered apical F-actin morphology, which stain positive for mucus, indicating MCC trans-differentiation into Goblet cells. Magnified MCCs are indicated by yellow boxes. **(B)** Quantification of total MCC numbers and proportion of PNA(+) MCCs between st. 32 and 46. **(C)** Notch inhibition by DAPT treatment reduces the number of MCCs showing signs of trans-differentiation (red arrowheads). Normal MCCs indicated by yellow arrowheads. **(D)** Quantification of total MCC numbers and number of MCCs showing trans-differentiation (TD) morphology. Mann Whitney test, ns *P* > 0.05 = not significant, * *P* < 0.05. **(E)** Inhibition of Jak1/2 signaling by Ruxolitinib (Ruxo.) or inhibition of Thyroid hormone signaling by Propylthiouracil (PTU) decreases the number of MCCs showing signs of trans-differentiation (TD, blue box) and increases the number of MCCs with apoptotic/shedding morphology (red boxes; cf. **Fig. S2 E-G**). Normal MCC indicated by yellow box. **(F)** Quantification of total MCC numbers (upper panel; Mann Whitney test, ns *P* > 0.05 = not significant) and proportions of normal (yellow), trans-differentiating (blue), apoptotic/shedding (red) MCC phenotypes (lower panel; χ^2^ test, *** *P* < 0.001).

MCC to Goblet cell trans-differentiation was proposed in conditions associated with persistent inflammation such as Asthma or COPD (Chronic obstructive pulmonary disease) (Tyner et al., 2006), but this idea remains controversial and the opposite conversion was proposed as well (Ruiz García et al., 2019). In the mammalian airways, Interleukin-13 (IL-13) leads to Janus kinase 1 (Jak1) activation that phosphorylates Signal transducer and activator of transcription 6 (STAT6) (Tyner et al., 2006). Activated STAT6 (STAT6-p) leads to expression of the transcription factor *Spdef*, which in turn activates *Mucin5AC* transcription (Park et al., 2007; Parker et al., 2013). At the same time, STAT6-p was shown to directly repress *Foxj1*, a transcription factor required for maintenance of motile cilia (Gomperts et al., 2007). The IL-13 induced effects are modulated by Epidermal growth factor (EGF) signaling, which enhances Jak1 activity and exhibits an anti-apoptotic effect (Tyner et al., 2006) (**Fig. S5A**). We also observed loss of epidermal *foxj1* expression in *Xenopus* (**Fig. S5B**), but IL-13 was unlikely the trigger, because the gene is missing from the frog genome. The IL-13 effects in human MCCs could be blocked by inhibition of Notch signaling, and STAT6/Notch were shown to synergistically promote Goblet cells in the mouse (Danahay et al., 2015; Guseh et al., 2009). Together these findings suggested that Notch activation could lead to similar effects in *Xenopus* as IL-13 in mammals.

To determine how Notch signaling could be activated at distance to the lateral line, we analyzed the expression of the Notch ligands *jag1*, *delta-like1* (*dll1*) and *delta-like4* (*dll4*) on histological sections. This showed transient upregulation of Notch ligand expression in the mesoderm underlying the epidermis in stages of MCC trans-differentiation (**Fig. S6A-C**). Additional analysis of *hes1* expression confirmed increased Notch signaling activation in the epidermis, but at lower levels as compared to cells directly overlaying the lateral line primordium (**Fig. S6D,E**). Therefore, we tested treatment of tadpoles with DAPT throughout later stages of development, which significantly reduced trans-differentiation (**Fig. 3C,D** and **Fig. S5D**).

Next, we tested the effects of Jak signaling on *Xenopus* MCCs through inhibition of Jak1/2 by Ruxolitinib. Ruxolitinib induced MCC shedding distant to the lateral line (**Fig. 3E,F** and **Fig. S5D**), demonstrating similar anti-apoptotic effects for Jak1/2 as in mammalian MCCs. We wondered how Jak/STAT signaling could be elevated systemically to account for the observed anti-apoptotic and trans-differentiation promoting effects. Thyroid hormone (TH) can potentiate STAT as well as EFGR signaling via the MAPK/ERK cascade (Cheng et al., 2010). The *Xenopus* thyroid primordium forms at st. 33 and matures into the thyroid gland by st. 43 (Fini et al., 2012), thus, coinciding with MCC trans-differentiation. Therefore, we blocked TH signaling by application of Propylthiouracil (PTU). The application of PTU during stages of MCC trans-differentiation induced apoptosis in MCCs (**Fig. 3E,F** and **Fig. S5D**), similar to Ruxolitinib. These findings indicated that TH signaling enhances STAT activity to reach levels sufficient to prevent Notch induced apoptosis, similar to the effect of EGF signaling in mammalian MCCs (**Fig. S5C**).

To study how MCC to Goblet cell trans-differentiation is executed at the sub-cellular level, we investigated ciliation, basal bodies and F-actin organization during this process through direct comparison of trans-differentiating and normal MCCs within the same specimens. This revealed a decrease in acetylation of MCC cilia (Ac.-α-tubulin), loss of normal basal body spacing (Centrin4-GFP/CFP) and reduced Cep164-mCherry localization to basal bodies (**Fig. 4A** and **Fig. S7A,B**). Additionally, we observed drastic remodeling of the apical and sub-apical F-actin network in trans-differentiating MCCs, which resulted in an altered cell morphology and loss of polarized basal body alignment visualized by RFP-Clamp (**Fig. S7B,C**).

**Fig. 4.**
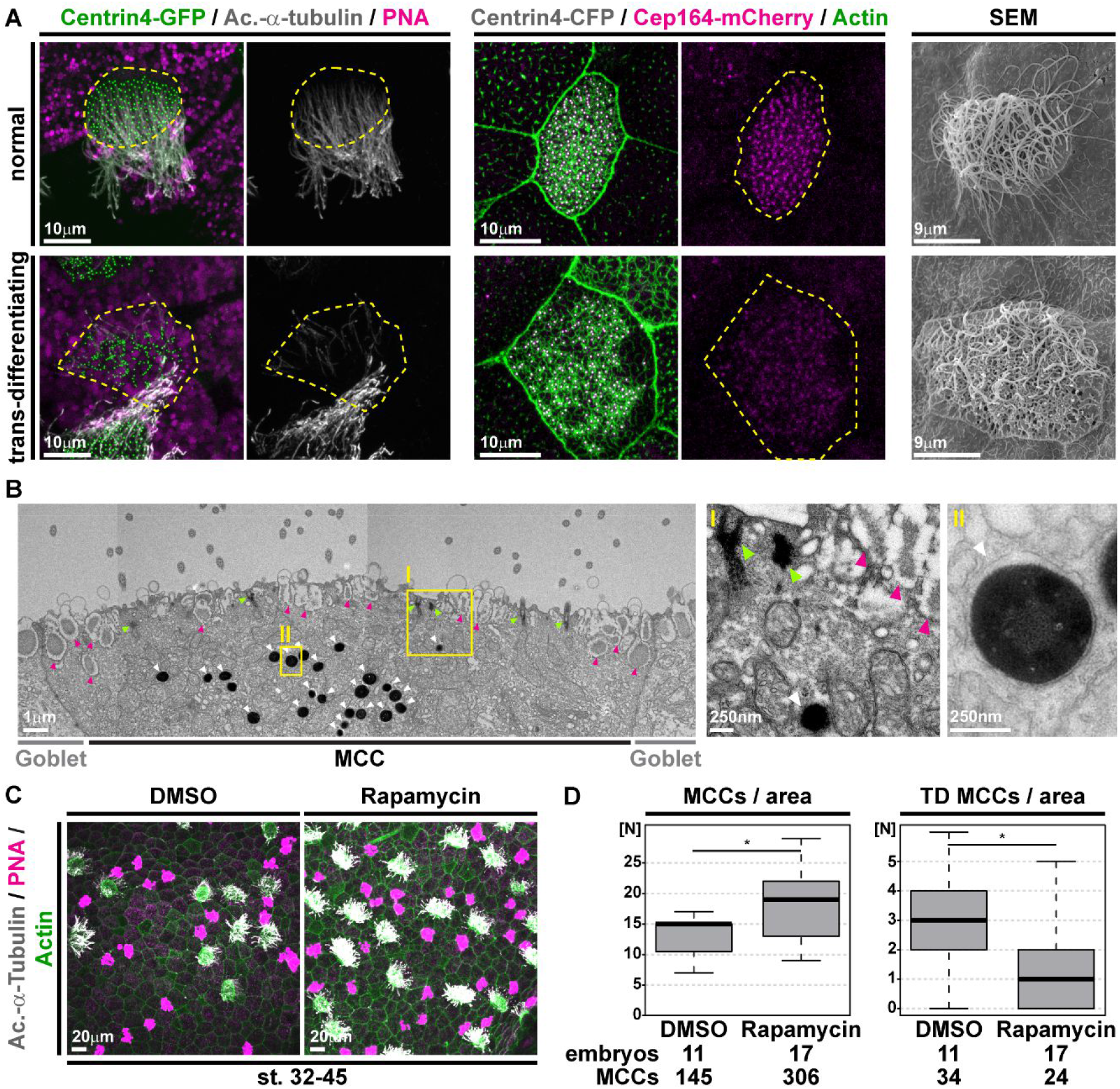
Inhibition of mTOR signaling prevents cilia retraction, basal body degradation and MCC loss. **(A)** Confocal and scanning electron microscopy (SEM) micrographs of normal and trans-differentiating MCCs reveal disorganized basal bodies (Centrin4-GFP, green / Centrin4-CFP, grey), cilia de-acetylation (Ac.-α-Tubulin, grey), F-actin remodeling (Actin, green) and loss of basal body distal appendages (Cep164-mCherry) during MCC cilia retraction. Cf. **Fig. S7 B** for quantifications. **(B)** Sagittal section and transmission electron microscopy (TEM) shows basal bodies (green arrowheads), mucus granules (magenta arrowheads) and an enrichment of lysosomes (white arrowheads) in a trans-differentiating MCC. Magnified areas are indicated by yellow boxes. Cf. **Fig. S8 C**. Image was reconstructed from multiple individual micrographs. **(C)** Confocal micrographs of the epidermis stained for MCCs (Ac.-α-Tubulin, grey), F-actin (Actin, green) and mucus (PNA staining, magenta) demonstrate an increase in MCCs and reduced trans-differentiation in Rapamycin treated tadpoles. **(D)** Quantification of total MCC numbers and number of MCCs showing trans-differentiation (TD) morphology. Mann Whitney test, * *P* < 0.05.

While Ac.-α-tubulin staining was strongly reduced or absent in trans-differentiating MCCs, SEM analysis showed persisting cilia with various degrees of length reduction (**Fig. 4A**), like during primary cilia retraction upon cell cycle re-entry (Izawa et al., 2015). We concluded that MCC cilia retraction followed a similar process as described for primary cilia, which was further supported by loss of expression of the cell cycle inhibitor p27 in trans-differentiating MCCs (**Fig. S7D,E**).

Next, we wondered what happens to the >100 basal bodies, which are a specific feature of MCCs. We visualized cilia and basal bodies in MCCs by mGFP and Centrin4-CFP, and analyzed tadpoles at st. 44/45. This revealed late stage trans-differentiating MCCs with few cilia and basal bodies, which localization was altered from the apical surface towards the deeper cytoplasm (**Fig. S8A**). In some cases, cytoplasmic Centrin4-CFP signals were associated with mGFP, suggesting basal body packaging into vesicles for degradation.

We followed-up on this finding by transmission electron microscopy and analyzed MCCs at various stages of trans-differentiation as judged by the amount of mucus in those cells (**Fig. S8B**). We observed undocked basal bodies, but none in cytoplasmic vesicles. Instead, we detected increased numbers of electron-dense structures reminiscent of lysosomes (Hurbain et al., 2017) as well as structurally incomplete basal bodies (**Fig. 4B** and **Fig. S8B-D**). We confirmed that trans-differentiating MCCs were enriched for lysosomes by Lysotracker staining in mRFP-injected specimens (**Fig. S9A,B**). Together, these data indicated active degradation of ciliary and basal body material in trans-differentiating MCCs.

To visualize possible Centrin4-CFP uptake into early lysosomes, we co-injected mRNA encoding human Lamp1-mCherry (Lamp1-mCherry) and analyzed MCCs at st. 43-46. We found that MCCs were enriched for Lamp1-mCherry-vesicles, confirming our Lysotracker results (**Fig. S9C**). We were not able to detect hLamp1-mCherry(+) trans-differentiating MCCs, while non-targeted MCCs were trans-differentiating at st. 43/44 (**Fig. S9C**), and ciliation persisted until st. 46 (**Fig. S9D**). These findings indicated that Lamp1-mCherry overexpression delayed trans-differentiation.

Impaired lysosome function upon Lamp1 overexpression was recently demonstrated in human pancreatic cancer cells (Chen et al., 2019), and lysosomal stress was reported to trigger autophagy, similar to nutrient deprivation (Huber and Teis, 2016; Pan et al., 2019), indicating that upregulated autophagy could prevent MCC trans-differentiation. Thus, we treated tadpoles with Rapamycin to stimulate autophagy through specific inhibition of the mTOR complex 1 (mTORC1). Rapamycin treatment during stages of trans-differentiation and analysis of MCCs at st. 45 revealed significantly reduced MCC loss and a reduction of MCCs showing features of trans-differentiation (**Fig. 4C,D**). This supported our hypothesis that the trans-differentiation process is inhibited by upregulation of autophagy.

## Discussion

We demonstrate that *Xenopus* epidermal MCCs can be lost by apoptosis or by trans-differentiation, two processes induced by Notch signaling. Trans-differentiation occurs in MCCs that are protected from apoptosis by Thyroid-dependent elevation of Jak/STAT signaling (**Fig. S5C**). MCC to Goblet cell trans-differentiation involves cilia de-acetylation and shortening, similar to primary cilia retraction, as well as F-actin remodeling and degradation of basal bodies. These processes can be inhibited by Rapamycin, which suggests a similar positive role for autophagy in MCC ciliation as described for primary cilia (Wang et al., 2015). Primary cilia formation is induced upon serum starvation, which inhibits mTORC1 and promotes autophagy (Boukhalfa et al., 2019). How precisely autophagy modulates cilia formation, length and maintenance remains incompletely resolved. Nevertheless, emerging evidence suggests that the balance between autophagy, endocytosis and proteasomal degradation processes influences cilia formation and resorption as well as maintenance of centrioles and basal bodies (Boukhalfa et al., 2019; Izawa et al., 2015; Liang et al., 2016; Paridaen et al., 2013).

Our results reveal deep evolutionary conservation of signaling and cellular mechanisms between mucociliary epithelia of the mammalian lung and the ciliated epidermis of basal fish and amphibia. We also confirm the proposal that MCCs can trans-differentiate into secretory cells, which are indistinguishable from other Goblet cells by our analyses, demonstrating a complete direct change of cell identity. By examining the process of apoptosis and trans-differentiation, we provide insights into the molecular basis of tissue remodeling in chronic lung disease. Importantly, the roles of autophagy and the use of Rapamycin have been considered in the context of COPD and Asthma, were tested in various models, but with variable effects (Kennedy and Pennypacker, 2016; Lam et al., 2013; Mitani et al., 2016; Mushaben et al., 2011; Wang et al., 2018). Cell type-specific effects of Rapamycin in the airways were discussed, but without reaching a consensus. mTOR signaling was also implicated in the potentiation of JAK/STAT activity (Bartalucci et al., 2017; Yokogami et al., 2000), suggesting additional beneficial effects for Rapamycin in the prevention of IL-13 induced trans-differentiation in the airways. With respect to MCC trans-differentiation, inactivation of the MCC-specific transcriptional program suppresses p27 and releases the cell-cycle block characteristic of ciliated cells (Izawa et al., 2015). As Goblet cells continue to divide after specification (cf. **Fig. 1A**), it is possible that former MCCs could re-enter the cell cycle after trans-differentiation and contribute to excessive Goblet cell hyperplasia in chronic lung disease (Hogan et al., 2014; Tyner et al., 2006). Thus, our findings provide important information that might serve as basis to facilitate more tailored tests of Rapamycin intervention in specific cases of chronic lung disease.

## Acknowledgments

We thank: S. Schefold, J. Groth and S. Kayser for expert technical help; Walz and Driever labs for sharing resources; S. Arnold, A. Classen, L. Davidson, W. Driever, R. Harland, G. Pyrowolakis, A. Schambony, J. Wallingford for discussions and/or critical reading; Xenbase (RRID:SCR_004337), NXR (RRID:SCR_013731) and EXRC for Xenopus resources; Light Imaging Center Freiburg and EM Core Facility BIOSS for microscope use.

## Author contributions

AT, MB, MHA, PW: Xenopus experiments; MHE: Electron microscopy; AT, PW: experimental design, planning and analysis; PW: study design and supervision, coordinating collaborative work, manuscript preparation.

## Materials and Methods

### Animals

Wild-type *Xenopus laevis* were obtained from the European *Xenopus* Resource Centre (EXRC) at University of Portsmouth, School of Biological Sciences, UK. Frog maintenance and care was conducted according to standard procedures and based on recommendations provided by the international *Xenopus* community resource centers NXR and EXRC as well as by Xenbase (http://xenbase.org). This work was done in compliance with German animal protection laws and was approved under Registrier-Nr. G-18/76 by the state of Baden-Württemberg. All experiments were conducted in embryos derived from at least two different females and independent *in vitro* fertilizations.

### Fertilization and microinjection of *Xenopus* embryos

*X. laevis* eggs were collected and *in vitro*-fertilized with wild-type or transgenic sperm, then cultured and/or microinjected by standard procedures (Sive et al., 2000). *In vitro* fertilization with p27∷GFP [*Xla.Tg(Xtr.cdknx:GFP)^Papal^*, RRID:EXRC_0043] (Carruthers et al., 2003) frozen sperm was performed by NXR protocol. Frozen sperm was removed from liquid nitrogen and immediately swirled in a 37°C water bath for 30sec, then resuspended in 500μl room-temperature 1/3x Modified Frog Ringer’s solution (MR) by using a micropestle. Sperm mixture was then added to the clutch of eggs and mixed into the monolayer for 3-5min, then flooded with 1/3x MR. After fertilization, embryos were injected with mRNAs and DNAs at the four-cell stage using a PicoSpritzer setup in 1/3x MR with 2.5% Ficoll PM 400 (GE Healthcare, #17-0300-50), and were transferred after injection into 1/3x MR containing 50μg/ml Gentamycin. Drop size was calibrated to about 7–8nl per injection.

### mRNAs and DNA constructs used in this study

mRNAs encoding membrane-RFP/GFP (gift from the Harland lab), Centrin4-GFP/CFP (Antoniades et al., 2014; Park et al., 2008), Clamp-RFP (Park et al., 2008), Cep164-mCherry (Tu et al., 2018), hLamp1-mCherry (this study) were prepared using the Ambion mMessage Machine kit using Sp6 (#AM1340) supplemented with RNAse Inhibitor (Promega #N251B), and diluted to 30–80ng/μl for injection into embryos. hLamp1-mCherry was generated by subcloning hLamp1-mCherry from pcDNA3.1 (Addgene #45147) to pCS107 using primers BamH1-hLamp1mCherry-F *(3‘-AAAAAAGGATCCATGGCGGCCCCCGGCAGCGC-5‘)* and hLamp1mCherry-EcoR1-R *(3‘-AAAAAAGAATTCTTACTTGTACAGCTCGTCCATG-5‘). The* sequence was verified by Sanger sequencing. The α-tub∷GFP (Chung et al., 2014) plasmid was injected at 5-10ng/μl after purification using the PureYield Midiprep kit (Promega, #A2492).

### Drug treatments of *Xenopus* embryos

Drug treatment of embryos started/was terminated at the indicated stages. Embryos were incubated in 24-well plates containing 2ml of 1/3MR + Gentamycin. DMSO (Roth #A994.2) was used as vehicle control at the same volume as specific drugs where indicated. Media were replaced every day. All treatments were carried out at roomtemperature (RT) and protected from light. DAPT (N-[(3,5-Difluorophenyl)acetyl]-L-al-anyl-2-phenyl]glycine-1,1-dimethylethyl ester; Sigma-Aldrich #05942) was dissolved in DMSO and used at 100-200μM (Myers et al., 2014). PTU (Propylthiouracil; Sigma-Aldrich #1578000; 10mM stock stored at −80°C for long term and −20°C for short term) was dissolved in Ultrapure water (ThermoFisher #10977035) and used at 1mM (Veenendaal et al., 2013). Rapamycin (Sigma-Aldrich #37094) was dissolved in DMSO and used at 100nM. Ruxolitinib (Selleckchem #S1378) was dissolved in DMSO and used at 10μM.

### LysoTracker use in *Xenopus* embryos

For LysoTracker (Invitrogen #L7526 Green DND-26) treatment, embryos were incubated in 12-well plates containing 4ml of 1/3x MR and 50μg/mL of Gentamycin. 50μM of LysoTracker was added to the medium for 5min in the dark at RT. Embryos were washed 3x 3min in 1/3x MR and finally anesthetized in buffered MS222 (Sigma-Aldrich #E10521) solution during mounting and live cell imaging. Non-LysoTracker controls were not incubated with the compound and anesthetized in buffered MS222 solution during mounting and live cell imaging using the same microscope settings.

### Immunofluorescence staining and sample preparation

Whole *Xenopus* embryos, were fixed at indicated stages in 4% Paraformaldehyde (PFA) in PBS at 4°C overnight or 2h at RT, then washed 3x 15min with PBS, 2x 30min in PBST (0.1% Triton X-100 in PBS), and were blocked in PBST-CAS (90% PBS containing 0.1% Triton X-100, 10% CAS Blocking; ThermoFischer #00-8120) for 1h at RT. All antibodies were applied in 100% CAS Blocking over night at 4°C or 2h at RT (for secondary antibodies). Primary antibodies used: mouse monoclonal anti-Acetylated-α−tubulin (1:1000; Sigma/Merck #T6793), anti-α-tubulin (1:1000; Abcam #ab7291). Secondary antibodies used: AlexaFluor 405-labeled goat anti-mouse antibody (1:500; ThermoFisher #A31553). Actin was stained by incubation (30-120min at RT) with AlexaFluor 488- or 647-labeled Phalloidin (1:40 in PBST; Molecular Probes #A12379 and #A22287), mucus-like compounds in *Xenopus* were stained by incubation (overnight at 4°C) with AlexaFluor 647-labeled PNA (1:1000 in PBST; Molecular Probes #L32460). Detailed protocol was published in (Walentek, 2018).

### Whole mount *in situ* hybridization

Embryos were fixed in MEMFA (100mM MOPS pH7.4, 2mM EGTA, 1mM MgSO4, 3.7% (v/v) Formaldehyde) overnight at 4°C and stored in 100% Ethanol at −20°C until used. DNAs were purified using the PureYield Midiprep kit and were linearized before *in vitro* synthesis of anti-sense RNA probes using T7 or Sp6 polymerase (Promega, #P2077 and #P108G), RNAse inhibitor and dig-labeled rNTPs (Roche, #3359247910 and 11277057001). Embryos were *in situ* hybridized according to (Harland and Biology, 1991), bleached (Sive et al., 2000) after staining and imaged. Sections were made after embedding in gelatin-albumin with Glutaraldehyde at 50-70μm as described in (Walentek et al., 2012).

### TUNEL assay

Embryos were fixed in MEMFA overnight at 4°C and stored in 100% Ethanol at −20°C until use. Embryos were bleached before staining. TUNEL staining was performed according to respective manufacturer protocol using Terminal Deoxynucleotidyl Transferase kits (Invitrogen #10533065 or Roche #03333574001), dig-UTP (Roche, #3359247910), anti-Digoxigenin AP antibody (1:3000; Roche #11093274910), and NBT/BCIP (Roche #11681451001). Samples were stopped in PBS, fixed briefly with 4% PFA in PBS and imaged.

### Sample preparation for electron microscopy

Sample primary fixation was done by 4% PFA, 2% Glutaraldehyde (Carl Roth #4157) in 0.1M Cacodylate Buffer (Science Services #11650). For transmission electron microscopy (TEM), the tissue was then post-fixed in 0.5% osmium tetroxide (Science Services #E19150) in ddH_2_O for 60min on ice and then washed 6x in ddH_2_O. The tissue was incubated in 1% aqueous uranyl acetate solution (Science Services #E22400-1) for 2h in the dark and washed 2x in ddH_2_O. Dehydration was performed by 15min incubation steps in 30%, 50%, 70%, 90% and 2x 100% Ethanol (Fisher Scientific #32205) and 2x 100% Aceton (Sigma-Aldrich #179124). After embedding in Durcupan resin (Sigma-Aldrich #44611 and #44612), ultrathin sections (55nm) were performed using a UC7 Ultramicrotome (Leica), collected on Formvar-coated (Science Services #E15830-25) copper grids (Plano #G2500C). Post-staining was done for 1min with 3% Lead Citrate (Delta Microscopies #11300). For scanning electron microscopy (SEM), the fixated samples were dehydrated in 70%, 80%, 90% and 100% Ethanol (each step for 1h at RT) and incubated in a 1:1 solution of Ethanol and Hexamethyldisilazan (HMDS; Carl Roth #3840.2) for 30min. After incubation in 100% HMDS, the solvent was allowed to evaporate. The dehydrated tissue was mounted onto sample holders and sputtered with gold using a Polaron Cool Sputter Coater E 5100.

### Light imaging, electron microscopy and image processing

All confocal imaging was performed using a Zeiss LSM880 and Zeiss Zen Black software. Whole embryo bright-field or fluorescent images were done on a Zeiss AxioZoom setup and Zeiss Zen Pro Blue software. Sections were imaged on a AxioZoom or AxioImager.Z1 microscope and Zeiss Zen Pro Blue software. Fluorescence images were processed in ImageJ/Fiji (maximum intensity projections, reconstruction of tile scans, selection of indicated planes, brightness/contrast adjustment, merging of channels) (Schindelin et al., 2012). Brightfield images were adjusted for color balance, brightness and contrast using Adobe Photoshop. TEM imaging was done using a Zeiss Leo 912 transmission electron microscope. SEM imaging was done using a scanning electron microscope (Leo 1450 VP scanning). Electron microscopy including TEM (transmission electron microscopy) and SEM (scanning electron microscopy) images were adjusted for brightness and contrast, and pseudo-colored using Adobe Photoshop.

### Analysis of trans-differentiated MCCs (ciliation, Actin network, basal body integrity)

Imaging was performed using the same settings within individual experiments on embryos which were injected with equal amounts of mRNAs. Images were processed using ImageJ to adjust brightness/contrast and to generate maximum intensity projections. For quantification of ciliation rates, images from similar areas were acquired (cf. **Fig. S5 D**). Trans-differentiated MCCs were designated by presence of mucins and/or impaired apical actin network and impaired ciliation. For quantification of cilia acetylation, basal body polarization and presence of Cep164-mCherry, normal and trans-differentiated MCCs from the same embryos were analyzed. For presentation of some representative examples, deep optical planes from the F-actin channel containing muscle fibers were manipulated to remove muscle signals.

### Statistical evaluation

Stacked bar graphs were generated in Microsoft Excel, boxplots were generated in R (the line represents the median; 50% of values are represented by the box; 95% of values are represented within whiskers; values beyond 95% are depicted as outliers). Statistical evaluation of experimental data was performed using Wilcoxon sum of ranks (Mann-Whitney) test (https://astatsa.com/WilcoxonTest/), or χ^2^ test (http://www.physics.csbsju.edu/stats/t-test.html) as indicated in figure legends. Sample sizes for all experiments were chosen based on previous experience and used embryos derived from at least two different females. No randomization or blinding was applied.

## Supplementary Material

Supplementary Figures S1-S9.

