## Supplementary Figures S1-S9 for "A changing signaling environment induces multiciliated cell trans-differentiation during developmental remodeling"

Tasca et al. Fig. S1

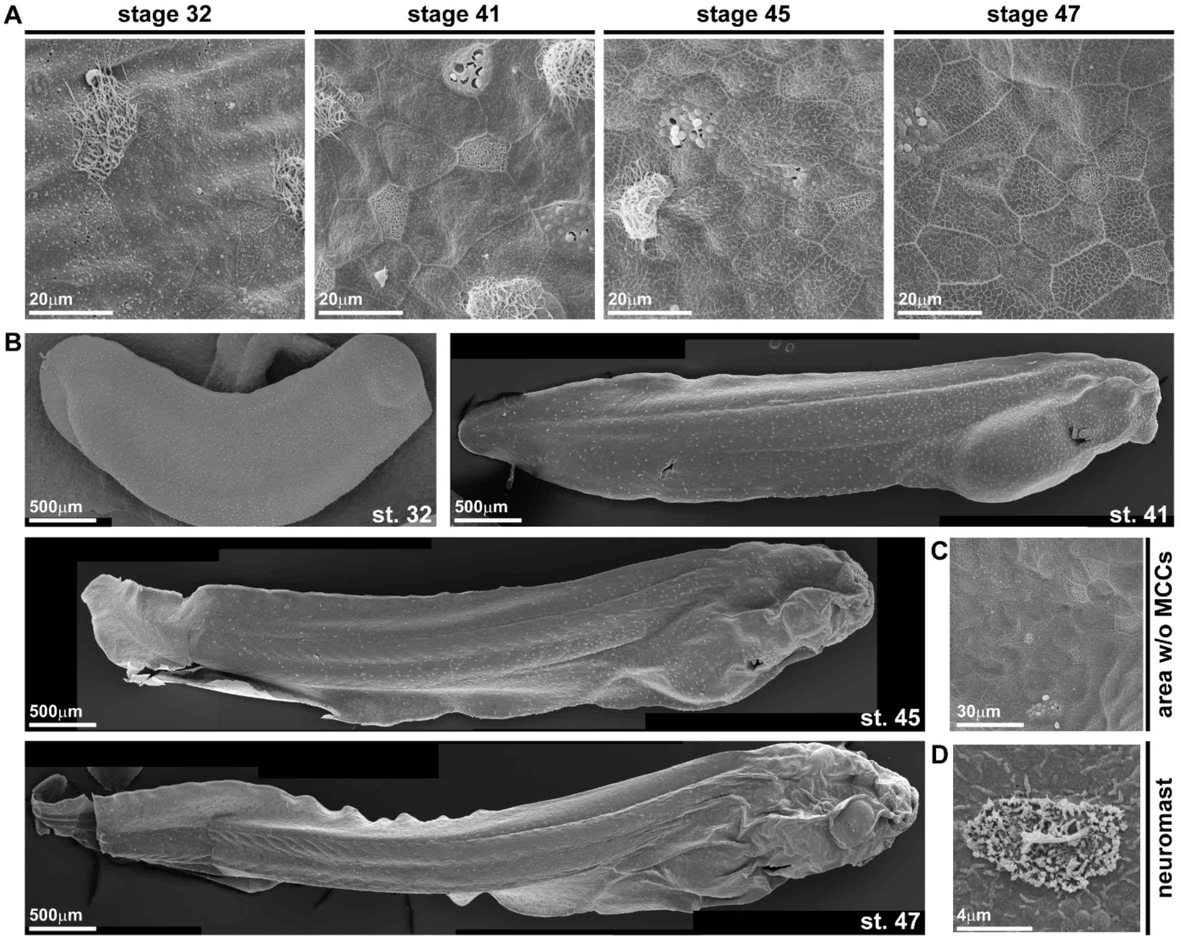


### Fig. S1.

Original micrographs used for characterization of epidermal MCC loss during *Xenopus* development in Fig. 1. **(A-D)** Non pseudo-colored scanning electron micrographs from developmental stages 32 through 47. Cf. Fig. 1. **(A)** Analysis of the changing composition of mucociliary cell types in the epidermis shows progressive loss of MCCs, while ionocytes, small secretory cells and mucus-secreting Goblet cells remain present. **(B)** Analysis of MCC-loss patterns on whole tadpoles. Images were reconstructed from multiple individual micrographs. **(C)** Magnification of skin area devoid of MCCs at stage (st.) 45 reveals presence of lateral line neuromast. **(D)** Magnified image of a neuromast.

**Tasca et al. Fig. S2**

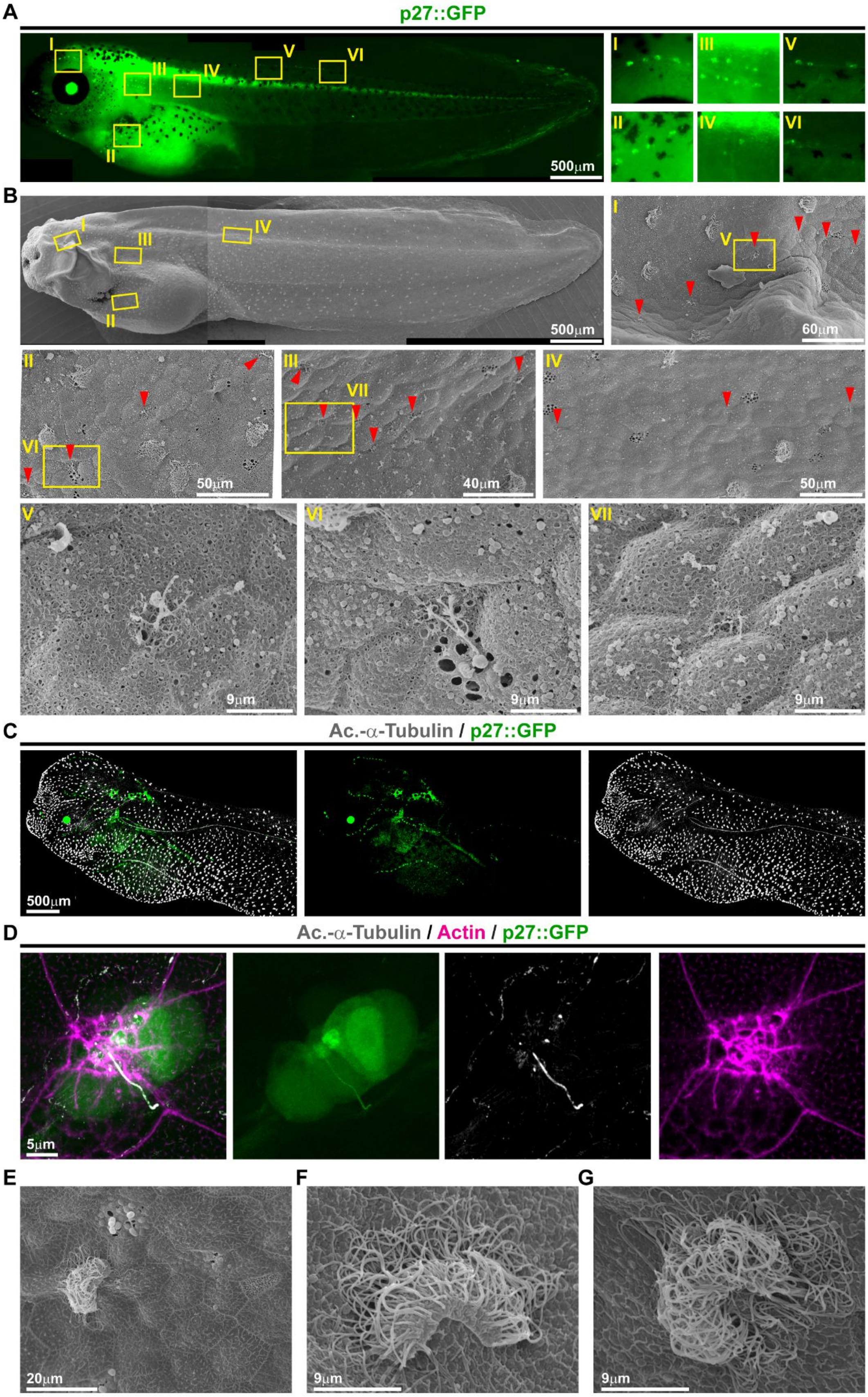

**Fig. S2.**

Local loss of MCCs coincides with the emergence of the lateral line. **(A,B)** Correlative light and electron microscopy (CLEM) analysis of p27 promotor-driven GFP (p27::GFP) transgenic tadpole identifies GFP expression in *Xenopus* neuromasts of the lateral line system. Images were reconstructed from multiple individual micrographs. **(A)** Fluorescent light microscopy image depicts GFP expression. Yellow boxes indicate location of magnified images. **(B)** Scanning electron micrographs of the same tadpole depicted in A confirm presence of neuromasts (red arrowheads) in locations of GFP expression. Yellow boxes indicate location of magnified images. **(C)** Confocal micrograph of p27::GFP transgenic tadpole that was stained with antibody to visualize MCCs (Ac.-$\alpha$-Tubulin, grey). **(D)** Confocal micrograph of p27::GFP transgenic tadpole that was stained with antibody to visualize cilia (Ac.-$\alpha$-Tubulin, grey) and phalloidin-647 to visualize F-actin (Actin, magenta) shows GFP expression in a subset of ciliated neuromast cells. **(E-G)** Scanning electron micrographs of areas with reduced ciliation reveal MCCs with abnormal morphology, indicating shedding of MCCs from the epithelium. Images in A-C were reconstructed from multiple individual micrographs.

Tasca et al. Fig. S3

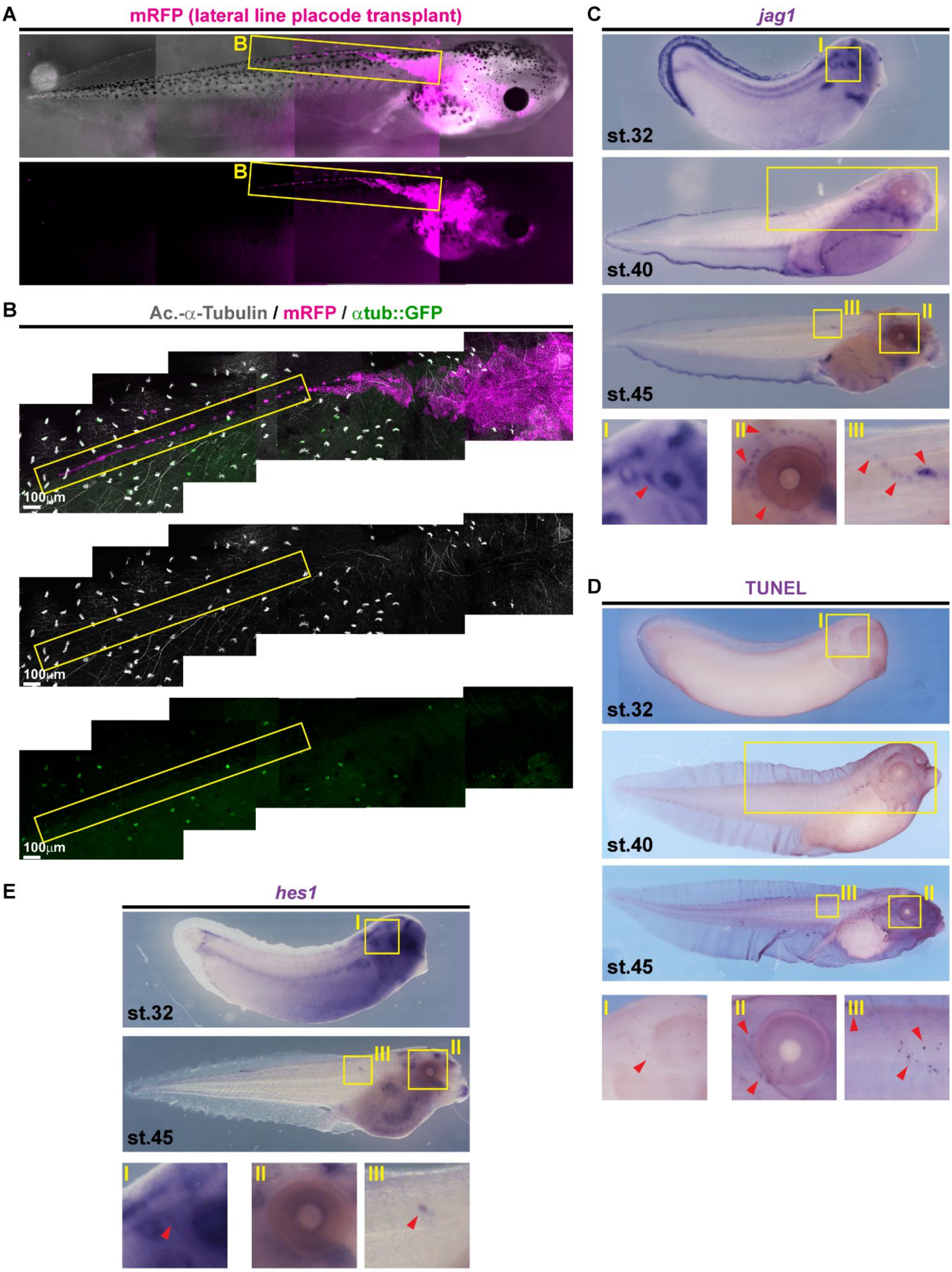

**Fig. S3.**

Local loss of MCCs correlates with neuromast emergence, Notch ligand expression and induction of apoptosis. **(A,B)** Micrographs of transplanted lateral line primordium and neuromasts (marked by membrane-RFP, mRFP) reveal loss of MCCs (marked by α-tubulin promotor-driven GFP [αtub::GFP, green] and Acetylated-α-Tubulin [Ac.-α-Tubulin, grey] antibody staining) in areas where neuromasts have emerged. Images were reconstructed from multiple individual micrographs. **(A)** Fluorescent light microscopy image depicts whole tadpole at stage 45 which received a mRFP-labeled transplant of a lateral line primordium at st. 18. Yellow boxes indicate location used for confocal micrograph in B. N = 2. **(B)** Confocal micrograph of area depicted in B. Yellow boxes indicate the image area depicted in Fig. 2 A. **(C)** *In situ* hybridization shows *jagged1* (*jag1*, purple) expression in the lateral line primordium and in neuromasts at stages 32, 40 and 45. Yellow boxes indicate location of magnified images. Cf. Fig. 2 C. **(D)** TUNEL staining (purple) reveals apoptotic cells clustering along the lateral line migration paths at stages 32, 40 and 45. Yellow boxes indicate location of magnified images. Cf. Fig. 2 D. **(E)** *In situ* hybridization shows transient *hes1* (purple) expression in the lateral line primordium at stages 32 and 45. Yellow boxes indicate location of magnified images.

Tasca et al. Fig. S4

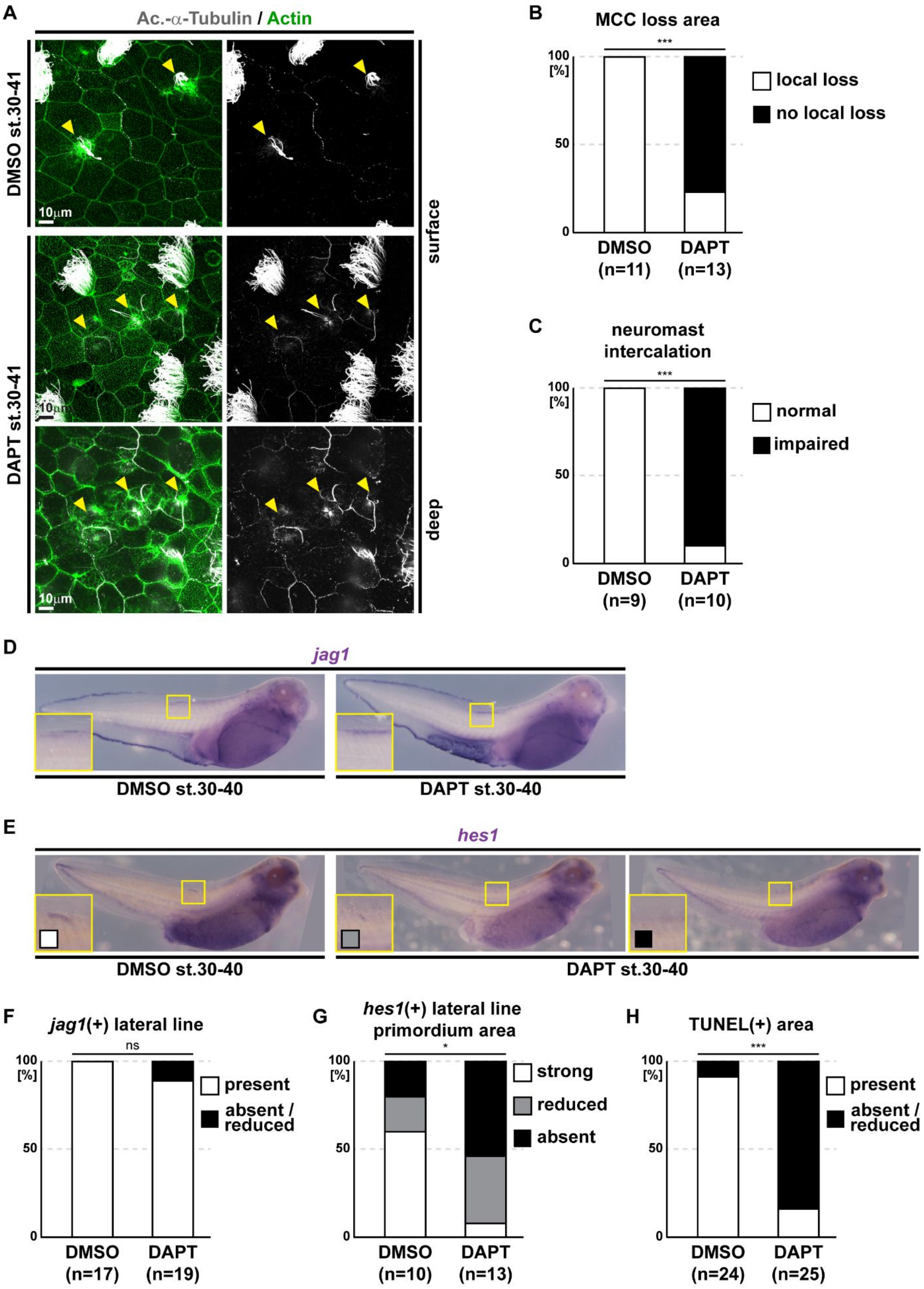

**Fig. S4.**

Notch signaling inhibition prevents MCC apoptosis, epidermal neuromast intercalation, but not lateral line migration and *jag1* expression. **(A)** Notch signaling inhibition by DAPT application prevents lateral line induced MCC (Ac.-$\alpha$-Tubulin, grey) loss and neuromast (yellow arrowheads) intercalation. F-actin (Actin, green) was used as counterstain in confocal images. Please also note impaired neuromast spacing in DAPT treated tadpoles. **(B)** Quantification of local MCC loss in control (DMSO) and DAPT treated tadpoles. $\chi^2$ test, *** $P < 0.001$. **(C)** Quantification of neuromast intercalation in control (DMSO) and DAPT treated tadpoles. $\chi^2$ test, *** $P < 0.001$. **(D)** *In situ* hybridization shows unaffected *jag1* (purple) expression in the lateral line primordium and in neuromasts in control (DMSO) and DAPT treated tadpoles. Yellow boxes indicate location of magnified images. **(E)** *In situ* hybridization shows reduced *hes1* (purple) expression in the lateral line primordium and the trunk in DAPT treated tadpoles as compared to controls (DMSO). Yellow boxes indicate location of magnified images. **(F)** Quantification of *jag1* expression in control (DMSO) and DAPT treated tadpoles. $\chi^2$ test, $P > 0.05$ = not significant. **(G)** Quantification of *hes1* expression in control (DMSO) and DAPT treated tadpoles. $\chi^2$ test, * $P < 0.05$. **(H)** Quantification of lateral line induced apoptosis (TUNEL staining, purple) in control (DMSO) and DAPT treated tadpoles. $\chi^2$ test, *** $P < 0.001$.

Tasca et al. Fig. S5

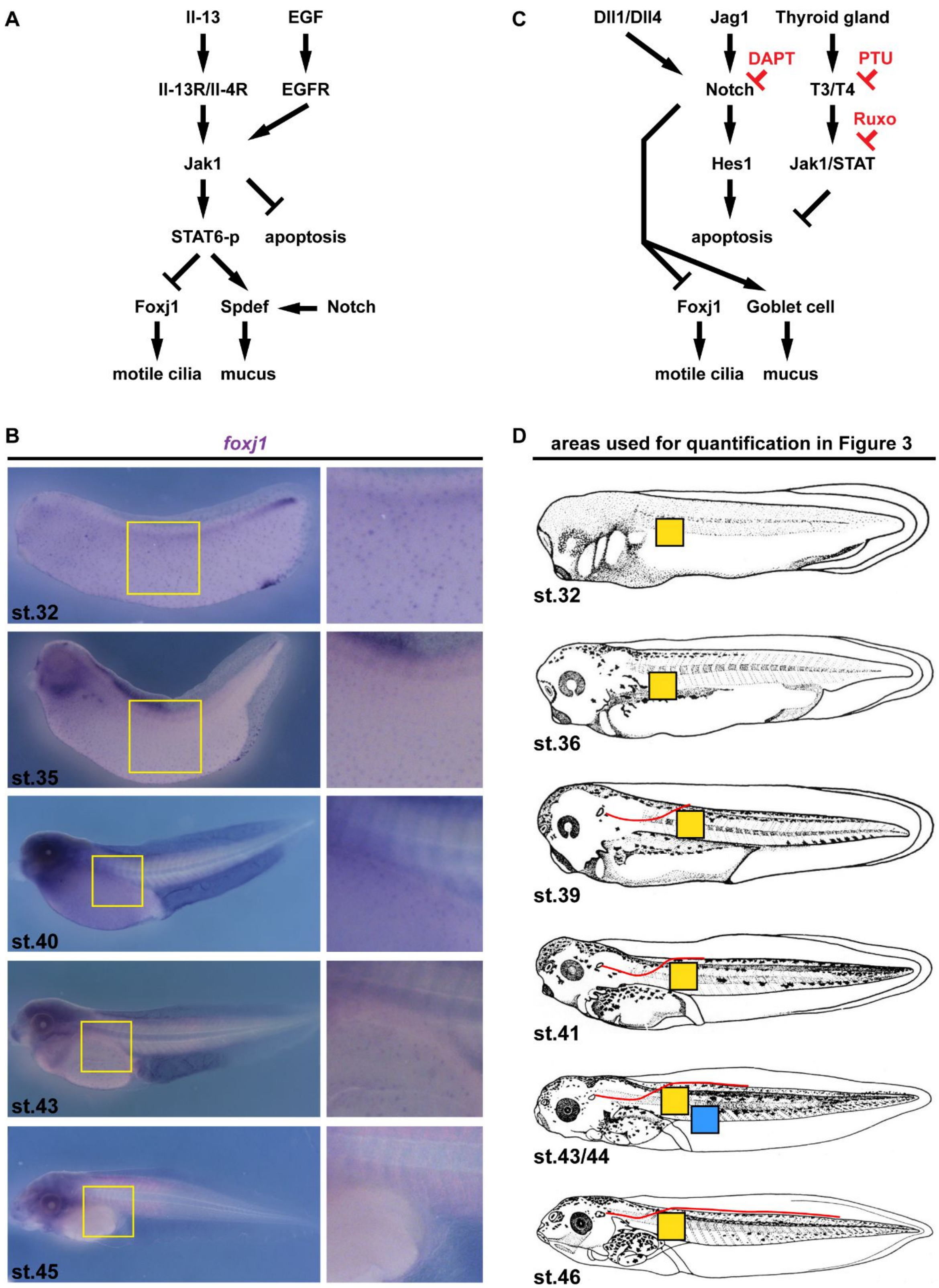

**Fig. S5.**

Global loss of MCCs through trans-differentiation into Goblet cells is regulated by Notch, Jak/STAT and Thyroid hormone signaling. **(A)** Schematic summary of signaling events described in the context of mammalian airway MCC to Goblet cell trans-differentiation. **(B)** *In situ* hybridization shows progressive loss of *foxj1* (purple) expression in the epidermis of *Xenopus* tadpoles during stages (32 to 45) of MCC trans-differentiation. Yellow boxes indicate location of magnified images. **(C)** Schematic summary of signaling events that lead to MCC apoptosis and trans-differentiation described in this study. The targets of pharmacological intervention as well as the small molecules used in this study are depicted in red. Ruxo = Ruxolitinib. **(D)** Schematic depiction of epidermal areas used for the analysis and quantification of MCCs in Fig. 3. Yellow boxes indicate areas used for MCC quantification in Fig. 3 A,B. Blue box indicates area used for MCC quantification in Fig. 3 C-F. Location of the dorsal lateral line is indicated in red.

Tasca et al. Fig. S6

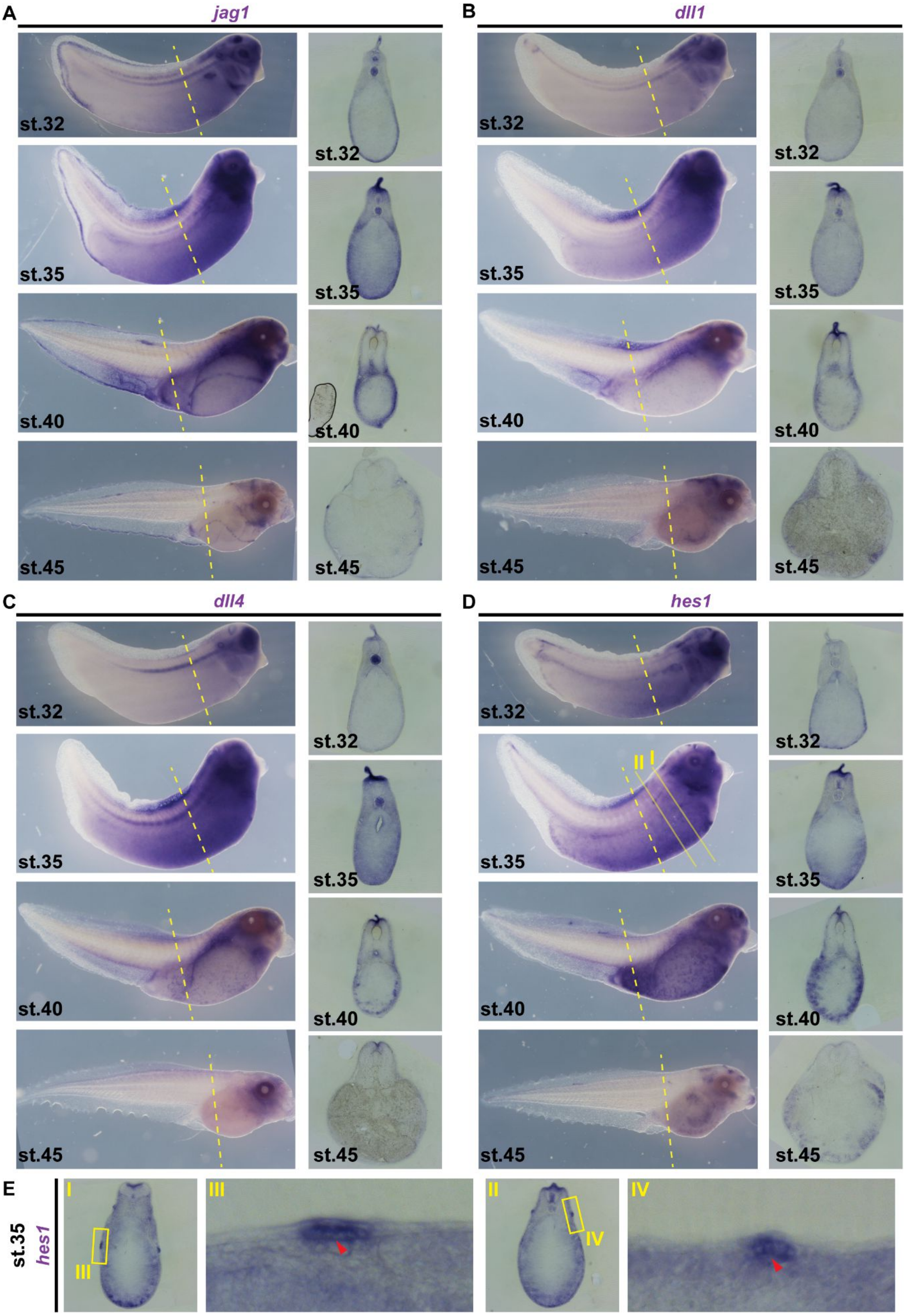

**Fig. S6.**

Expression analysis of Notch ligands *jag1*, *dll1* and *dll4*, and the Notch target *hes1*. **(A-D)** Whole mount *in situ* hybridization was conducted on specimens from the same batches for all four genes in stages 32 to 45. Histological sections reveal distribution of signals among various tissues. **(A-C)** The expression of the Notch ligands *jag1*, *dll1* and *dll4* in mesodermal tissues underlying the epidermis increases between st. 32 and st. 35, is maintained at high levels at st. 40, and decreases by st. 45, i.e. after trans-differentiation of MCCs is largely completed. **(D)** The expression of the Notch target *hes1* in mesodermal tissues as well as in the epidermis increases between st. 32 and st. 35, is maintained at high levels at st. 40, and decreases by st. 45. **(E)** Additional sections from st. 35 tadpole depicted in D reveal *hes1* expression in the epidermis, but especially elevated levels in epidermal areas directly overlying the migrating lateral line primordium (red arrowheads). Planes of sections in A-D are indicated by yellow dashed lines. Planes of sections depicted in E are indicated in D by solid yellow lines.

Tasca et al. Fig. S7

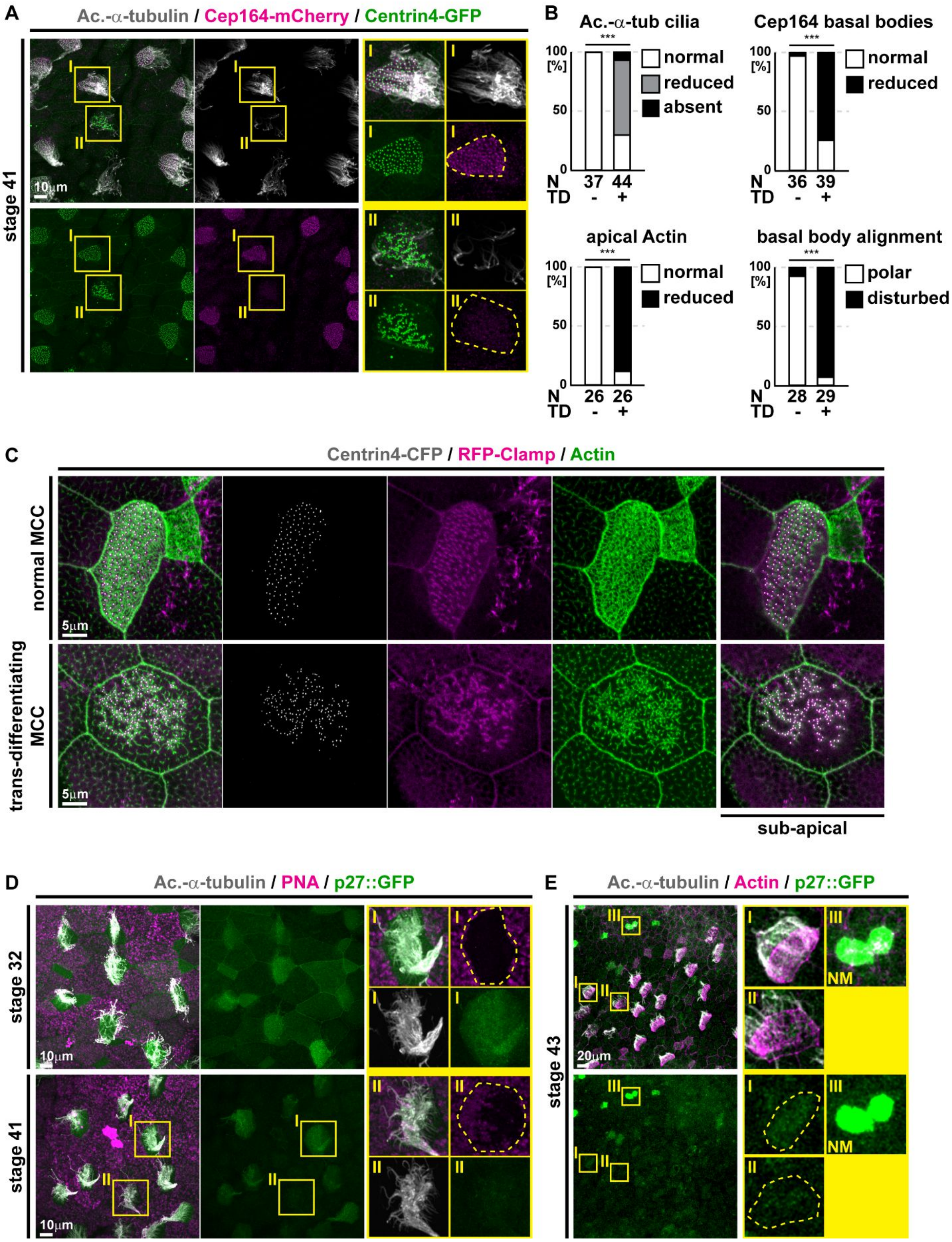

**Fig. S7.**

De-acetylation of cilia, F-actin remodeling, loss of basal body polarity, and de-repression of proliferation in trans-differentiating MCCs. **(A)** Confocal micrographs of normal and trans-differentiating MCCs reveal disorganized basal bodies (Centrin4-GFP, green), cilia de-acetylation (Ac.-$\alpha$-Tubulin, grey) and loss of basal body distal appendages (Cep164-mCherry) during MCC cilia retraction in the same specimen. Magnified areas are indicated by yellow boxes. **(B)** Quantification of MCC phenotypic differences between normal and trans-differentiating MCCs. N = number of MCCs. TD = trans-differentiating. $\chi^2$ test, *** $P < 0.001$. Cf. Fig. 4 A. **(C)** Confocal micrographs of a normal and a trans-differentiating MCC stained for basal bodies (Centrin4-CFP, gray), rootlets (RFP-Clamp, magenta) and F-actin (Actin, green) reveals loss of basal body alignment as well as apical and sub-apical F-actin organization. **(D)** Confocal micrographs of transgenic p27::GFP (green) expression in normal and trans-differentiating MCCs at stages 32 and 41. MCCs (Ac.-$\alpha$-Tubulin, grey), and mucus (PNA staining, magenta). Note the loss of GFP expression in MCCs with reduced cilia acetylation that start to express mucins as compared to normal MCCs within the same specimen. N = 3 embryos per stage. Magnified areas are indicated by yellow boxes. **(E)** Confocal micrograph of transgenic p27::GFP (green) expression in MCCs and Neuromasts (NM) at stages 43. MCCs (Ac.-$\alpha$-Tubulin, grey), and F-actin (Actin, magenta). Note the loss of GFP expression in MCCs with reduced cilia acetylation and altered apical Actin as compared to less trans-differentiated MCC and Neuromast within the same specimen. N = 5 embryos. Magnified areas are indicated by yellow boxes.

**Tasca et al. Fig. S8**

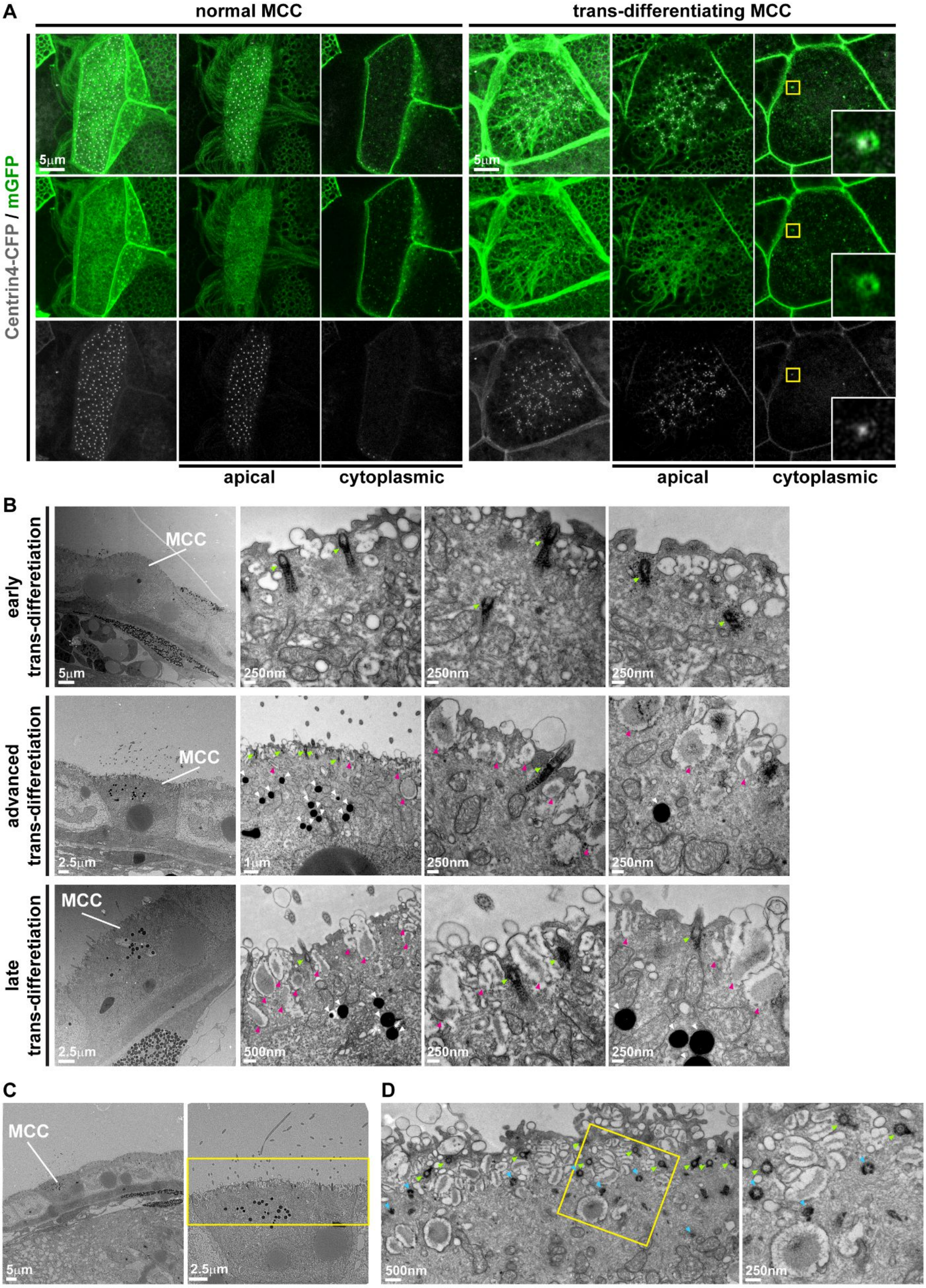

**Fig. S8.**

Degradation of basal bodies in trans-differentiating MCCs. **(A)** Confocal micrographs of normal and trans-differentiating MCCs reveal cytoplasmic basal bodies (Centrin4-CFP, gray) co-localizing with membrane vesicles (mGFP, green) in late stage trans-differentiating MCC. Magnified areas are indicated by yellow boxes. Note also the weakening of Centrin4-CFP signals per basal body in trans-differentiating MCC specimen. Normal MCCs, N = 7; trans-differentiating MCCs, N = 8; N = 3 embryos. **(B)** Sagittal sections and transmission electron microscopy (TEM) of MCCs with different degrees of trans-differentiation (number of mucus granules; magenta arrowheads) show basal bodies (green arrowheads), including some localizing to the cytoplasm. Additionally, advanced and late trans-differentiating MCCs are enriched for lysosomes (white arrowheads). Multiple magnified images are shown, which are each derived from different section planes of the same MCC depicted in the left panel. N = 9 MCCs from 1 embryo. **(C)** Sagittal sections and TEM of MCC shows enrichment of lysosomes in trans-differentiating MCC as compared to neighboring Goblet cells. The yellow box in the right panel, indicates MCC area depicted in Fig. 4 B. **(D)** Transversal section and TEM of trans-differentiating MCC shows intact basal bodies (green arrowheads) as well as structurally incomplete basal bodies (blue arrowheads). Magnified area is indicated by yellow box. N > 3 MCCs from 1 embryo.

Tasca et al. Fig. S9

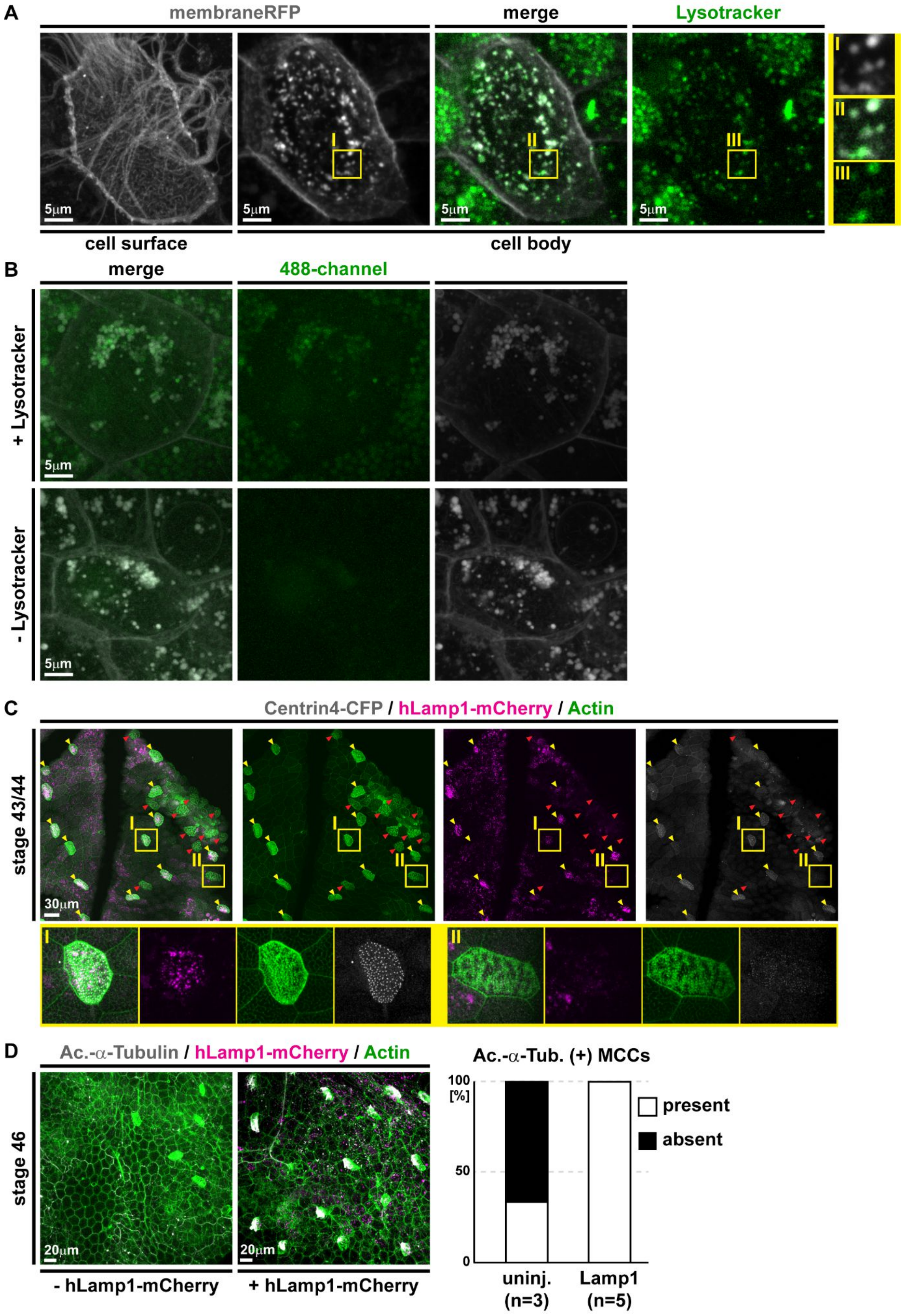

**Fig. S9.**

Tans-differentiating MCCs are enriched for lysosomes and Lamp1 overexpression interferes with trans-differentiation. **(A)** Confocal micrographs of trans-differentiating MCC reveal enrichment of cytoplasmic vesicles (mRFP, gray) that stain positive for Lysotracker (green). Magnified areas are indicated by yellow boxes. N = 33 MCCs / 11 embryos. **(B)** Comparison of 488-channel signals in Lysotracker treated (+Lysotracker) and non-treated (-Lysotracker) samples. Note also in A and B, that mucus granules in neighboring Goblet cells stain positive for Lysotracker, which is in line with mucus granule acidification required for Mucin packaging. **(C)** Confocal micrographs of normal and trans-differentiating MCCs within the same sample show enrichment of hLamp1-mCherry signals in targeted MCCs (yellow arrowheads) as compared to neighboring targeted Goblet cells. MCCs with hLamp1-mCherry overexpression do not show signs of trans-differentiation (yellow arrowheads) while trans-differentiation morphology is observed in non-targeted MCCs (red arrowheads). Sample was stained for Actin (F-actin, green). Magnified areas are indicated by yellow boxes. N = 48 MCCs / 4 embryos. Image was reconstructed from multiple individual micrographs. **(D)** Confocal micrographs of uninjected controls (- hLamp1-mCherry / uninj.) and hLamp1-mCherry injected (+ hLamp1-mCherry / Lamp1) specimens at st. 46. MCCs (Ac.-$\alpha$-Tubulin, grey), hLamp1-mCherry (magenta), F-actin (Actin, green). hLamp1-mCherry injected tadpoles present ciliated MCCs, while uninjected controls are devoid of cilia. Uninjected N = 3 embryos / 2 Ac.- $\alpha$-Tubulin (+) MCCs; hLamp1-mCherry injected N = 5 embryos / 61 Ac.- $\alpha$-Tubulin (+) MCCs.
